# Cluster Analysis for Protein Sequences

**DOI:** 10.1101/2025.03.14.643225

**Authors:** Md Rafsan Jani

## Abstract

This paper presents a comprehensive analysis of MMseqs2 clusters and traditional machine learning (ML) clustering algorithms, including KMeans and Hierarchical clusterings, in terms of protein sequences. The analyses are validated experimentally. The cluster analyses have been performed in the Astral Compendium protein sequences dataset hosted in the SCOPe database. The dataset is embedded using two pre-trained transformer models using Evolutionary Scale Modeling (ESM) to perform KMeans and Hierarchical clustering algorithms. Afterward, those four clusters are compared with `MMseqs2/Linclust` and `MMseqs2/easy-cluster` methods. After performing the experiment, `MMseqs2/Linclust` and `MMseqs2/easy-cluster` outperform traditional machine learning cluster algorithms by a considerable margin. This analysis demonstrates the superiority of the MMseqs2 clustering techniques over conventional machine learning clustering algorithms. The source code of the experiment is publicly available and readily accessible through: https://github.com/mrzResearchArena/protein-clustering.

## 1 Introduction

In metagenomics and metatranscriptomics studies, many novel species have been discovered in recent years which are not closely related to any organism. Consequently, those species are not well-annotated. Almost all proteins have structural similarities with others; sometimes, they share a common evolutionary origin [-1]. Furthermore, protein families can be distinguished by molecules that share significant sequence similarities [-2]. As an enormous number of proteins have structural similarities, we are required to minimize redundancy by employing clustering algorithms.

Protein clustering provides meaningful and persistent groupings of similar proteins. This information is required for functional annotation and efficient searching. In addition, protein clusters are collections of homologous proteins that carry almost similar functions and are associated with each other [-3]. Metagenomics datasets usually contain billions of protein sequences that expedite large-scale function annotation and structural prediction research [-4, -5]. Therefore, we need to categorize those enormous numbers of proteins that manifest similar structures and traits by employing a robust and effective clustering approach. In this paper, I investigate: whether protein sequences can be clustered employing typical ML clustering methods or alignment coverage-based clustering techniques are required, such as MMseqs2 clusters. I demonstrate which one is more convenient in terms of protein sequences via side-by-side comparisons.

## 2 Workflow

In this section, I elucidated my thought process on how I concluded the project. For instance: (1) What were my initial thoughts and the back of my mind? (2) How did I encounter issues? (3) How did I circumvent these pitfalls? (4) How was this experiment carried out?

In Figure 1, the workflow is illustrated that briefly depicts the entire working procedure.

**Figure 1.**
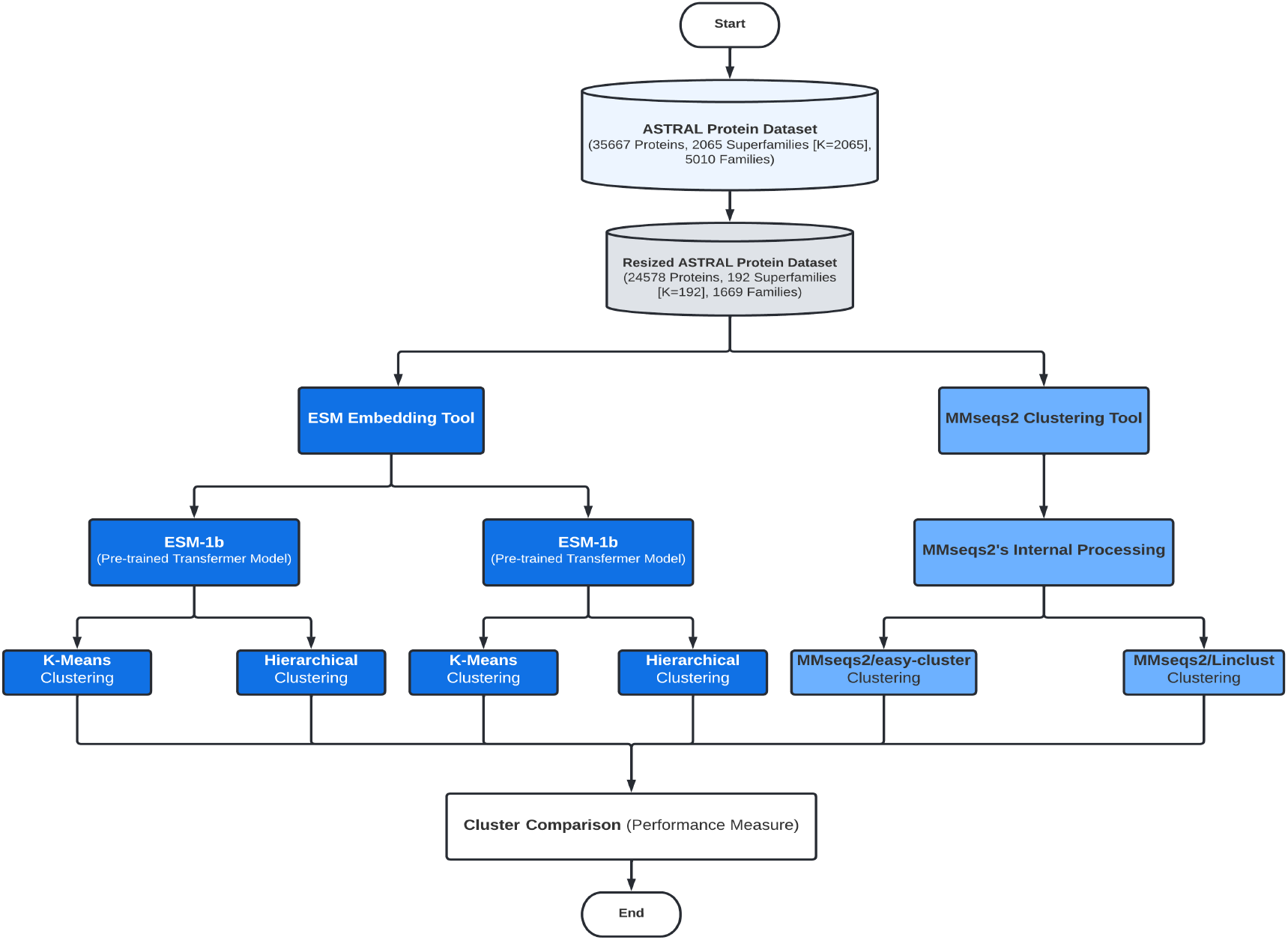
This image illustrates the entire working procedure.

**Step-1:** First and foremost, I tried to address the problem. Some of the issues are compiled in the back of my mind: (1) Why are clusters needed in terms of protein research? (2) What are the biological aspects behind it? (3) What is the significance of solving this particular problem? (4) How do MMseqs2 and ESM tools function? (5) Why do we need to compare traditional ML clustering methods and MMseqs2 clustering methods?

After having a general idea of the problem (though vaguely), I meticulously investigate the entire procedure step by step to understand the inside out of the project.

**Step-2:** Secondly, I studied what are the SCOPe and Astral Compendium databases; correlated their relationship by reading some of the papers [1, 6–8], and official documentation (https://scop.berkeley.edu/help). Subsequently, I became familiar with its file systems and understood how the database is organized (i.e., folds, superfamily, family) and the purposes of making the database. The explanation is given in Section 3.

**Step-3:** After understanding the database, I deliberately analyze the dataset and attempt to decipher new knowledge. For example: (1) how many superfamilies and families are included in my dataset, (2) how many unique organisms or creatures are present in my dataset, and so forth. The detailed findings from the dataset are given in Section 4.

**Step-4:** Afterward, I explore the ESM tool and understand how it embeds protein sequences by reading its papers [9]. I found that it has four core modules, namely: ESM-1b, ESM-1v, ESM-MSA-1, and ESM-IF1. Two of them, out of the four modules, are used as protein sequences embedding. To test the ESM tool, I embedded a few protein sequences and tried to cluster using KMeans and Hierarchical clusterings. The explanation is provided in Section 5 and Table 2.

**Step-5:** In this phase, I encountered a severe memory management issue while executing the conventional ML clustering algorithms, such as KMeans and Hierarchical clusterings, which are considered NP-hard algorithms. To alleviate this issue, the primary dataset was truncated. The details are provided in Section 4.

**Step-6:** After getting the resized protein sequences dataset, protein sequences have been embedded using both ESM-1b and ESM-1v pre-trained models. The explanation is provided in Section 5.

**Step-7:** After embedding protein sequences from ESM-1b and ESM-1v models, I ran two clusters for each model and scrutinized their performance by comparing them to the original labels of the dataset (i.e., which sequence belongs to which superfamily). The explanation is given in Sections 5 and 6.

**Step-8:** Thenceforth, I investigate the MMseqs2 tool by reading several papers [5, 4]. I noticed that the MMseqs2 tool is widely used in protein structure prediction, removing redundancy from sequences, sensitive sequence searching, and so forth; even the most advanced protein sequence predictor, AlphaFold2, used it to predict protein structure prediction in CASP14 competition [10]. The explanation is given in Section 5.

**Step-9:** Thereafter, I clustered my dataset using the `MMseqs2/Linclust` and the `MMseqs2/easy-cluster` algorithms and assessed the quality of clusters by comparing them to the labels of the original dataset. The explanation is given in Sections 5 and 6.

**Step-10:** Following this, the performances are compared to each other clusters. The explanation is given in Section 6.

**Step-11:** In the end, a comprehensive analysis is manifested of why these particular cluster algorithms, such as MMseqs2 clustering algorithms, outperform compared to conventional ML clustering methods. The details are given in Sections 7 and 8.

## 3 Dataset Description

In the experiment, a subset of the Astral Compendium proteins dataset is used from the SCOPe database (**S**tructural **C**lassification **o**f **P**roteins – **e**xtended). This database is hosted by UC Berkeley and Berkeley Lab. The aim of the SCOPe database is to provide authoritative information about proteins’ structural and evolutionary relationships [1, 6]. For instance, the SCOPe database gives information on the origin of a certain protein. Furthermore, this database is also conducive to identifying ancient homologous relationships, functional features, and so forth. In the SCOPe database, proteins are stored in a hierarchical order. (i.e., classes → folds → superfamilies → families → proteins → species) ^1^.

However, the Astral compendium datasets especially ensure non-redundant or low-redundancy sequences compared to other state-of-the-art databases, such as PDB and Pfam. Be that as it may, I am working on a set of domain sequences that are constructed according to the PDB SEQRES records with less than 95% sequence identity ^2^.

## 4 Data Analysis and Preprocessing

The subset of the Astral protein dataset comprises 35,667 protein sequences. In addition, after analyzing the dataset, 2065 unique superfamilies and 5010 unique families are identified ^3^. In the following dataset, protein sequences contain different organisms, such as humans (above eight thousand), mice (above two thousand), viruses (above one thousand), bacteria, fungi, and so forth ^4^.

In the following experiment, superfamilies are chosen above families for two key reasons. First and foremost, at each new release of the Astral dataset, all non-redundant sequences from each SCOPe superfamily are aligned using the MAFFT tool ^5^. Secondly, a Hidden Markov-based model is constructed to assure the quality of sequences by employing multiple sequence alignments for each superfamily [8, 2].

As the dataset contains 2065 unique superfamilies, I have unwittingly set the number of clusters to 2065 (K=2065) and embedded a minuscule number of proteins (2,500 out of 35,667) for testing clustering techniques. However, traditional machine learning clustering algorithms, by way of illustration, KMeans and Hierarchical clustering, are infeasible to execute due to their time and space complexities. The time complexity and space complexity of the KMeans algorithm are 𝒪 (*N ×K ×I*) and 𝒪 (*N ×* (*D* + *K*), respectively ^6^. Moreover, hierarchical clustering is more expensive compared to KMeans, whose time and space complexity are 𝒪 (*N* ^3^) and 𝒪 (*N* ^2^), respectively [11]. Thus, these algorithms consume enormous time even in a medium size dataset. Additionally, these clustering algorithms are considered an NP-hard problem. In traditional ML clustering algorithms, we can specify the number of clusters or groups.

In order to assure feasibility, I minimize the number of clusters in the dataset based on the minimum number of protein sequences containing each superfamily. As a matter of fact, the dataset has been slightly trimmed. The table 1 demonstrates the viability of selecting the cluster size (K denotes the number of clusters.).

**Table 1.**
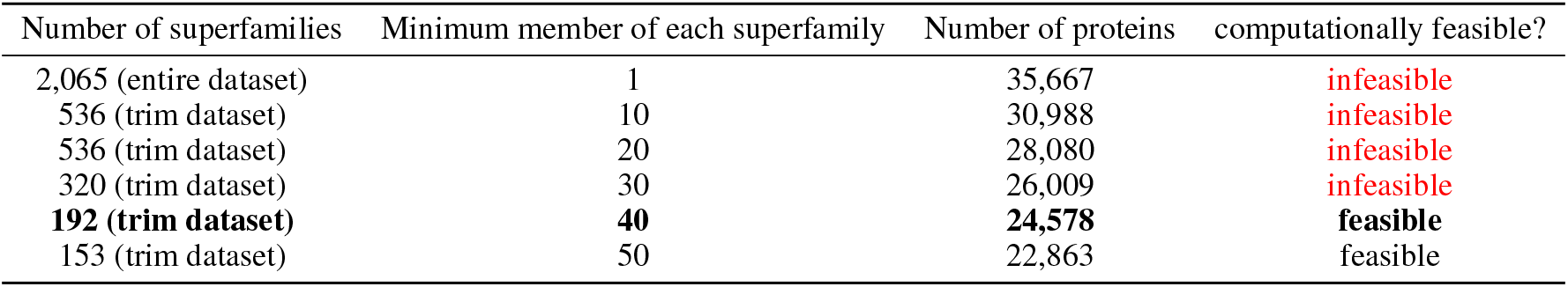
The table illustrates the viability of selecting the cluster size.

| Number of superfamilies | Minimum member of each superfamily | Number of proteins | computationally feasible? |
| --- | --- | --- | --- |
| 2,065 (entire dataset) | 1 | 35,667 | infeasible |
| 536 (trim dataset) | 10 | 30,988 | infeasible |
| 536 (trim dataset) | 20 | 28,080 | infeasible |
| 320 (trim dataset) | 30 | 26,009 | infeasible |
| **192 (trim dataset)** | **40** | **24,578** | **feasible** |
| 153 (trim dataset) | 50 | 22,863 | feasible |

Ultimately, I chose 192 superfamilies (K=192) instead of 2,065 because clustering the entire superfamily is computationally impractical within the limited resources. For this reason, I chose K=192 as the maximum number of proteins available within my computational resources; each superfamily contains at least 40 protein sequences. The red color indicates that these are the most expensive to compute (Table 1). However, those experiments are certainly possible if enough computer memory is available.

In Figure 2, the bar plots display the number of proteins belonging to each **superfamily**. In addition, in Figure 3, the bar plots display the number of proteins belonging to each **family** ^7^. The explanation behind the **family** exclusion over the **superfamily** is aforementioned in Section 4.

**Figure 2.**
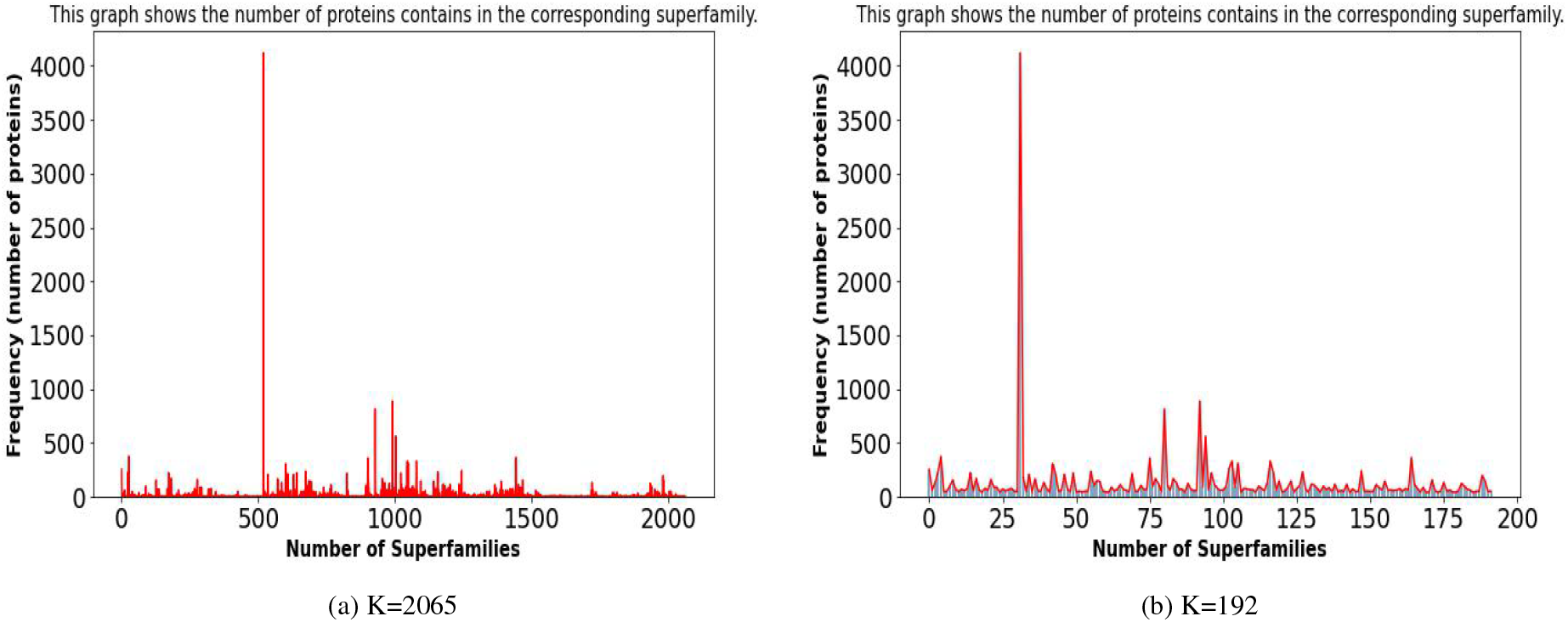
This bar graph illustrates the number of proteins found in each superfamily. In addition, image (a) K=2065 depicts the dataset before truncating, whereas image (b) K=192 depicts the dataset after trimming.

**Figure 3.**
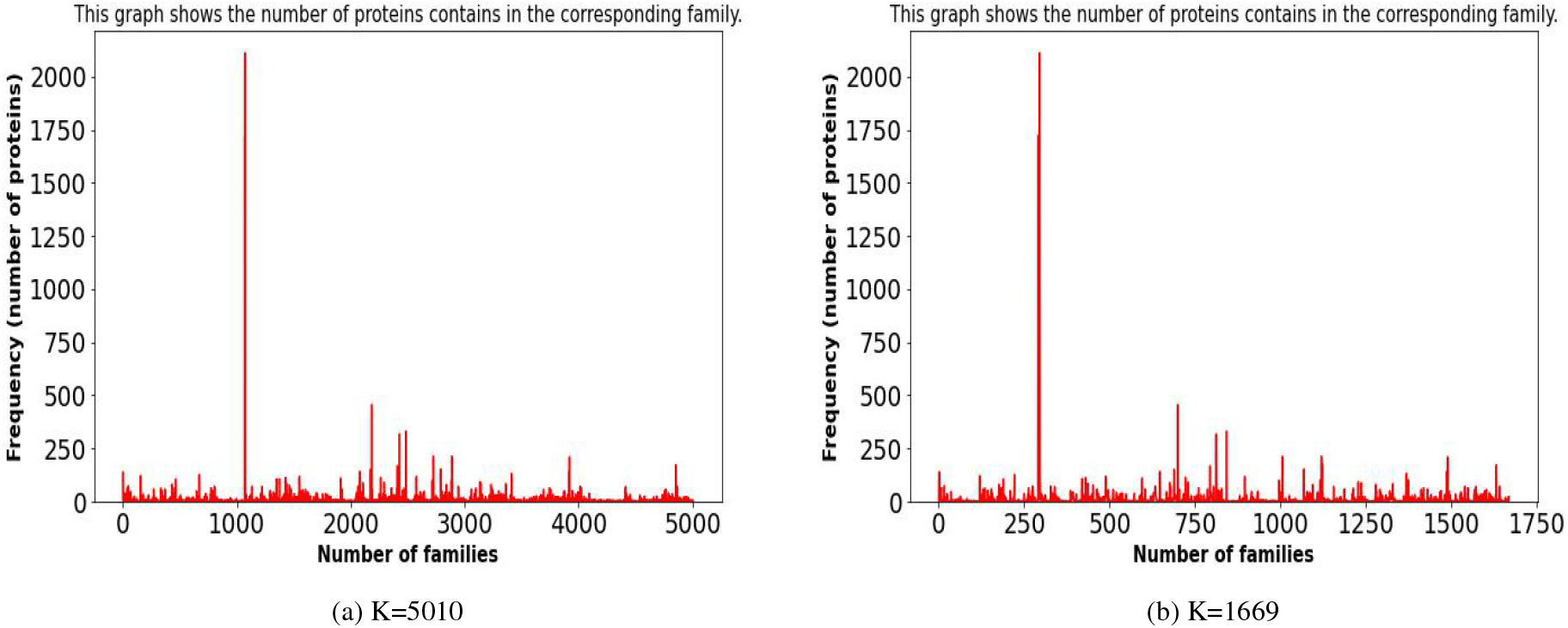
This bar graph illustrates the number of proteins found in each family. In addition, image (a) K=5010 depicts the dataset before truncating, whereas image (b) K=1669 depicts the dataset after trimming.

These images (Figure: 2 and Figure: 3) are atomic resolution and are challenging to interpret. Consequently, I have also generated photos in batches. Hence, we can visually investigate each **superfamily**. In Figure 4, only the first 15 superfamilies hosting the most members are displayed (i.e., protein sequences) ^8^.

**Figure 4.**
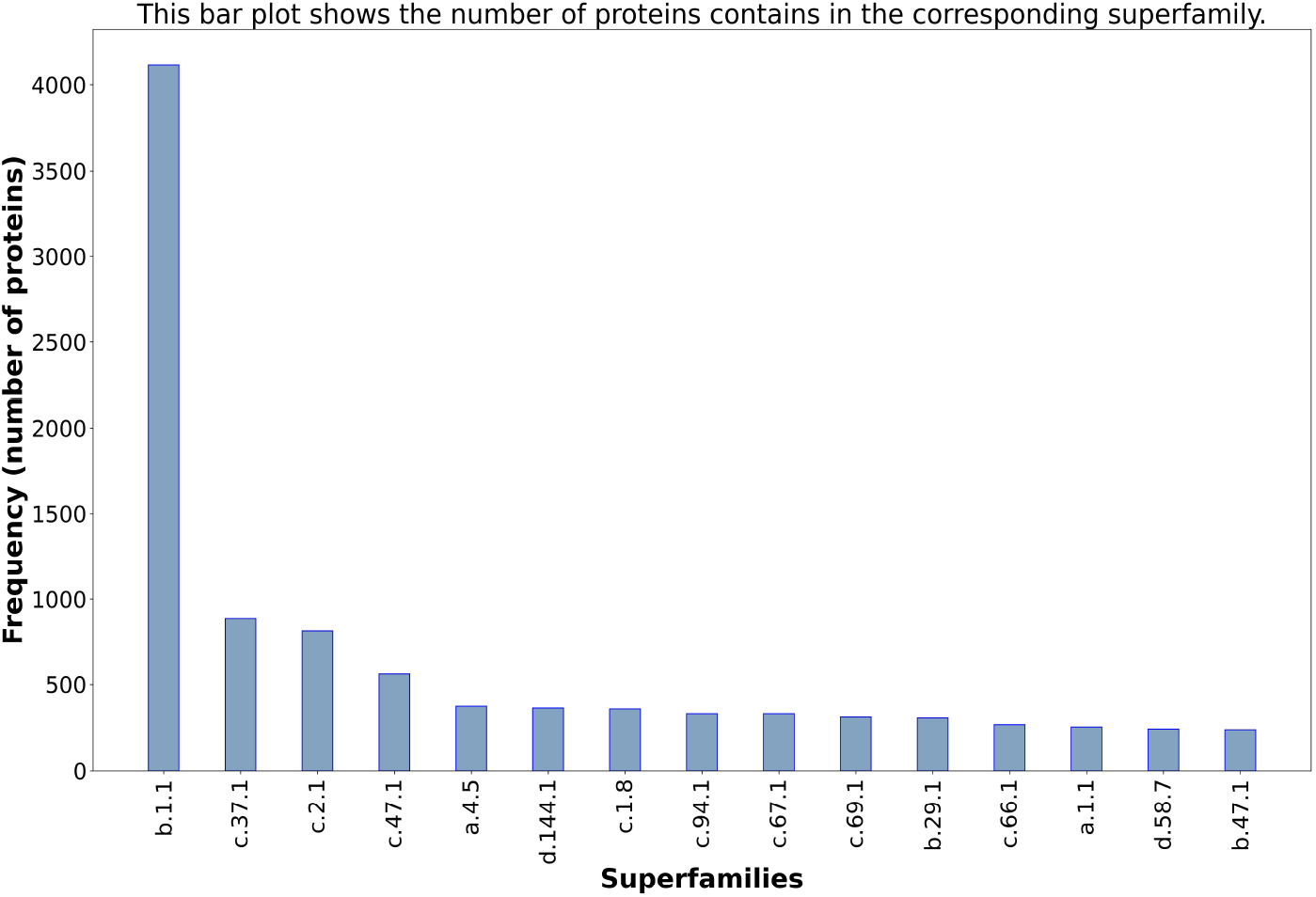
This bar graph illustrates the number of proteins found in the first 15 superfamilies that contains maximum protein sequences. However, this plot does not impact dataset truncating.

## 5 Experimental Validation

### 5.1 Preprocessing for Protein embedding

The ESM tool (Evolutionary Scale Modeling or Transformer Protein Language Models) supports a maximum of 1,022 residues for each protein. Despite the fact that ESM supports 1024 residues, two positions are reserved for beginning and terminating pointers. However, the size of residues in my dataset varies between 20 and 1664. This issue was resolved using the bigger sequences trimmed from the last regions of each sequence, and I kept 1022 residues from the beginning of each sequence ^9^.

### 5.2 Core Experiment

Afterward, I embedded the dataset employing two pre-trained transformer models, namely ESM-1b and ESM-1v ^10^. The embedding process is a bit tedious and has taken three and a half days as those pre-trained models require millions of parameters (i.e., weights and biases).

ESM has four pre-trained models for different purposes. However, two models among four are able to embed from raw protein sequences, including ESM-1b and ESM-1v. These two models share the same architecture. However, the difference between the two models is that ESM-1b was trained on the *UniProt50* database, whilst ESM-1v was trained on the *UniProt90* database. In addition, both ESM-1b and ESM-1v models are designed to predict structures, functions, variant effects, and other properties. Moreover, the ESM-1v has five different versions ^11^. In Table 2, the functionality of each pre-trained model (ESM) is briefly described.

**Table 2.** In this table, the information on pre-trained models (ESM) are briefly described.

| Transformer models | inputs | outputs | purposes |
| --- | --- | --- | --- |
| ESM-1b<br>(650 M parameters, 7.8 GB) | protein sequences | embedded protein sequences<br>(Output shape: number of sequences x 1280.) | unsupervised protein embedding to predict structures, functions, variant effects, and other properties |
| ESM-1v<br>(650M parameters, 7.8 GB) | protein sequences | embedded protein sequences<br>(Output shape: number of sequences x 1280.) | unsupervised protein embedding to predict structures, functions, variant effects, and other properties |
| ESM-IF1<br>(124 M parameters, 1.6 GB) | structures<br>(i.e., backbone atom coordinates as PDB and mmCIF formats) | protein sequences | inverse folding<br>(ProteinMPNN [12] can retrieve sequences approximately 52% accurately.) |
| ESM-MSA-1<br>(100 M parameters, 1.3 GB) | protein sequences<br>(length of sequences must be equal) | alignment matrices | alignments |

After getting two embedding metrics from ESM-1b and ESM-1v models, I performed two cluster algorithms: KMeans and Hierarchical clusterings for each embedding (Figure: 1). Furthermore, in both clusters, the *cosine similarity metric* has been used instead of the widely used Euclidean, Manhattan, and Minkowski distances ^12 13^. The main reason behind using the *cosine similarity metric* is that it considers the correlation between points. For instance, even though two proteins are far apart from each other (n-dimensional space), there is still a possibility that they are close together in terms of structural similarities. On the other hand, other distance metrics, such as Euclidean, Manhattan, and Minkowski, measure the distance between points without taking similarities into consideration. The cosine similarity score lies between 0 and 1. If the value is close to 1, it means two proteins are in the same orientation and have a high degree of similarities. In contrast, a score close to 0 indicates two proteins have fewer similarities. In addition, the *cosine similarity metric* is widely-used in Natural Language Processing (NLP) to determine the degree of similarity or dissimilarity between texts (i.e., plagiarism detection). The texts virtually look alike protein sequences. Therefore, the *cosine similarity metric* is unquestionably superior to other straightforward distance metrics, such as Euclidean and Manhattan.

After that, I explored the MMseqs2 tool, which is widely used to homology search and cluster enormous amounts of sequences, even billions of sequences. As MMseqs2 is faster than BLAST and reaches the same sensitivity as BLAST, many researchers are using this tool to expedite their speed in protein structure prediction. For instance, AlphaFold2 clustered to 30% sequence identity while enforcing a 90% alignment coverage of shorter sequences using MMseqs2/Linclust [10]. ColabFold incorporates MMseqs2 and gets 40-60 folds faster in homology search over groundbreaking AlphaFold2 structure prediction 13. OmegaFold harnesses the MMseqs2/Linclust cluster technique to reduce the redundancy, with sequence identity threshold of 100%, and keep the cluster centroids [14]. In the ESM, the transformer-based features are replaced by MMseqs2 features on the *UniClust90* dataset [9]. Additionally, the very recently released ESMFold (after performing my experiment) employs MMseqs2’s easy-cluster with 40% sequence identity (using default parameters). The performance is binned by the number of MMseqs2 hits while searching the training examples [15].

I conducted clusters using both `MMseqs2/Linclust` and `MMseqs2/easy-cluster` with 80% sequence identity and 5000 iterations. The hyper-parameters were tuned to get the optimal cluster centroids ^14^.

### 5.3 Experimental Setup

All the experiments were carried out using my personal computer and Google Colab Pro. My personal computer is equipped with 4-core processors (8-thread), each core having an AMD Ryzen Processor (7-3700U) with 2.3 GHz speed and 12 GB of memory. However, these configurations are inadequate due to insufficient memory for conventional ML clustering methods, such as KMeans and hierarchical clustering, as the time complexity is NP-hard. To alleviate the issue and expedite the working process, I subscribed to Google Colab Pro, which provides 25 GB of CPU memory and 15 GB of GPU memory.

## 6 Performance

In the end, the performances of clusters are compared to each other. After using `MMseqs2/Linclust` and `MMseqs2/easy-cluster`, I observed a mesmerizing result. Both `MMseqs2/Linclust` and `MMseqs2/easy-cluster` outperform compared to the traditional machine learning cluster algorithms, such as KMeans and hierarchical clustering, by a substantial margin. In addition, these two cluster algorithms show almost similar results. The cluster recovery rate is above 99%. On the other hand, the cluster recovery rate of typical ML algorithms lies between 40 to 65%. Upon assessing the performance, it was revealed that KMeans performed marginally better than the Hierarchical clustering algorithm.

However, the `MMseqs2/Linclust` provides approximately four times more cluster centroids compared to the `MMseqs2/easy-cluster`. Therefore, I would prefer the `MMseqs2/easy-cluster` technique, be that as it may, for the same reason, OmegaFold used the `MMseqs2/easy-cluster` method over `MMseqs2/Linclust` [14]. In the experiment, I kept both `MMseqs2/Linclust` and `MMseqs2/easy-cluster` to evaluate all the MMseqs2’s clustering techniques.

The cluster recovery rate is calculated using V-Measure, and the value lies between 0 to 1 [16]. V-Measure evaluates the quality of the clusters and ensures that the groups are virtually meaningful. However, various cluster assessment metrices, such as Rand Index, Silhouette Score, and Calinski-Harabasz Index, are widely used. In Table 3, a comparison between the clusters has been illustrated.

**Table 3.**
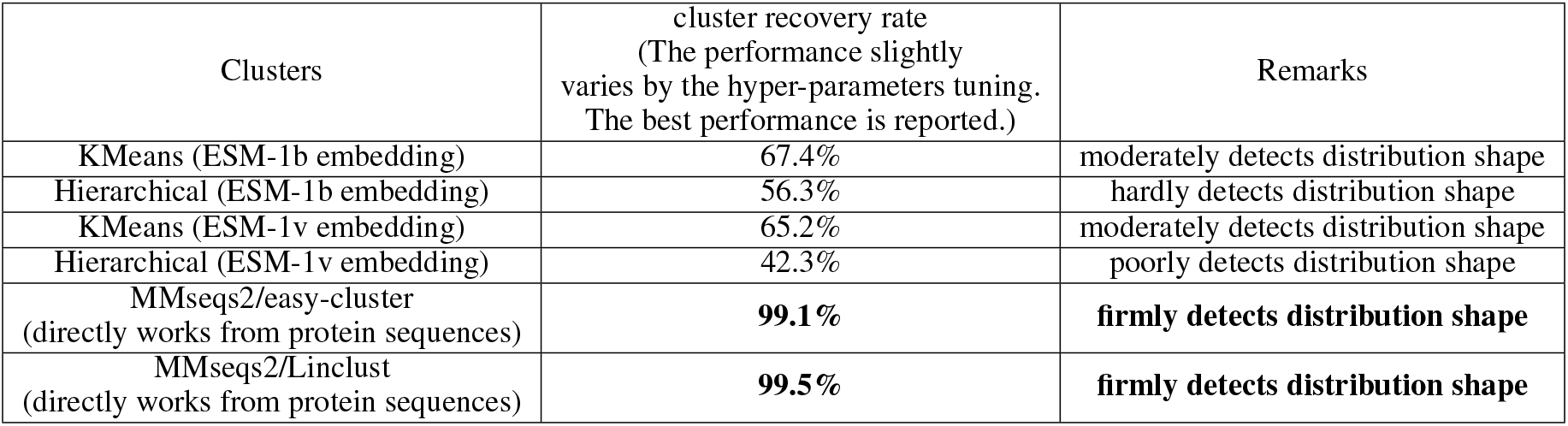
In this table, the cluster results are provided and compared to each other.

In Figure 5, I depict the detection result employing `MMseqs2/easy-cluster` algorithm because other traditional machine clusters poorly detect the distribution of superfamilies. However, `MMseqs2/Linclust` found almost the same distribution shape, like `MMseqs2/easy-cluster`. Hence, the distribution shape for the `MMseqs2/easy-cluster` and `MMseqs2/Linclust` methods is almost similar ^15^.

**Figure 5.**
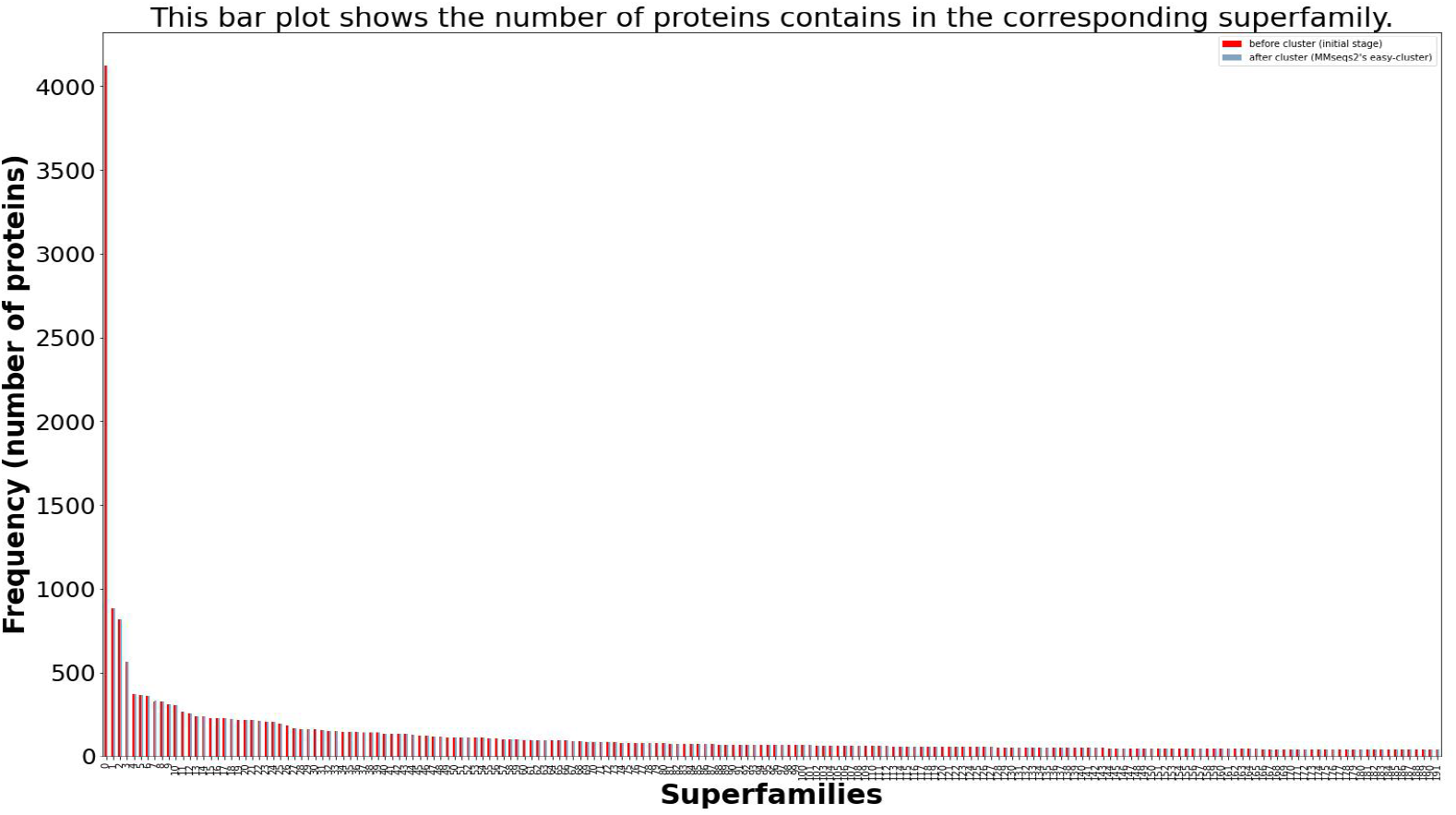
This image illustrates performance that is attained using MMseqs2 cluster algorithms.

In Figure 6, I show the first 15 maximum superfamilies, which contain maximum members (to see all comparisons ^16^) because it is easy to visualize compared to Figure 5.

**Figure 6.**
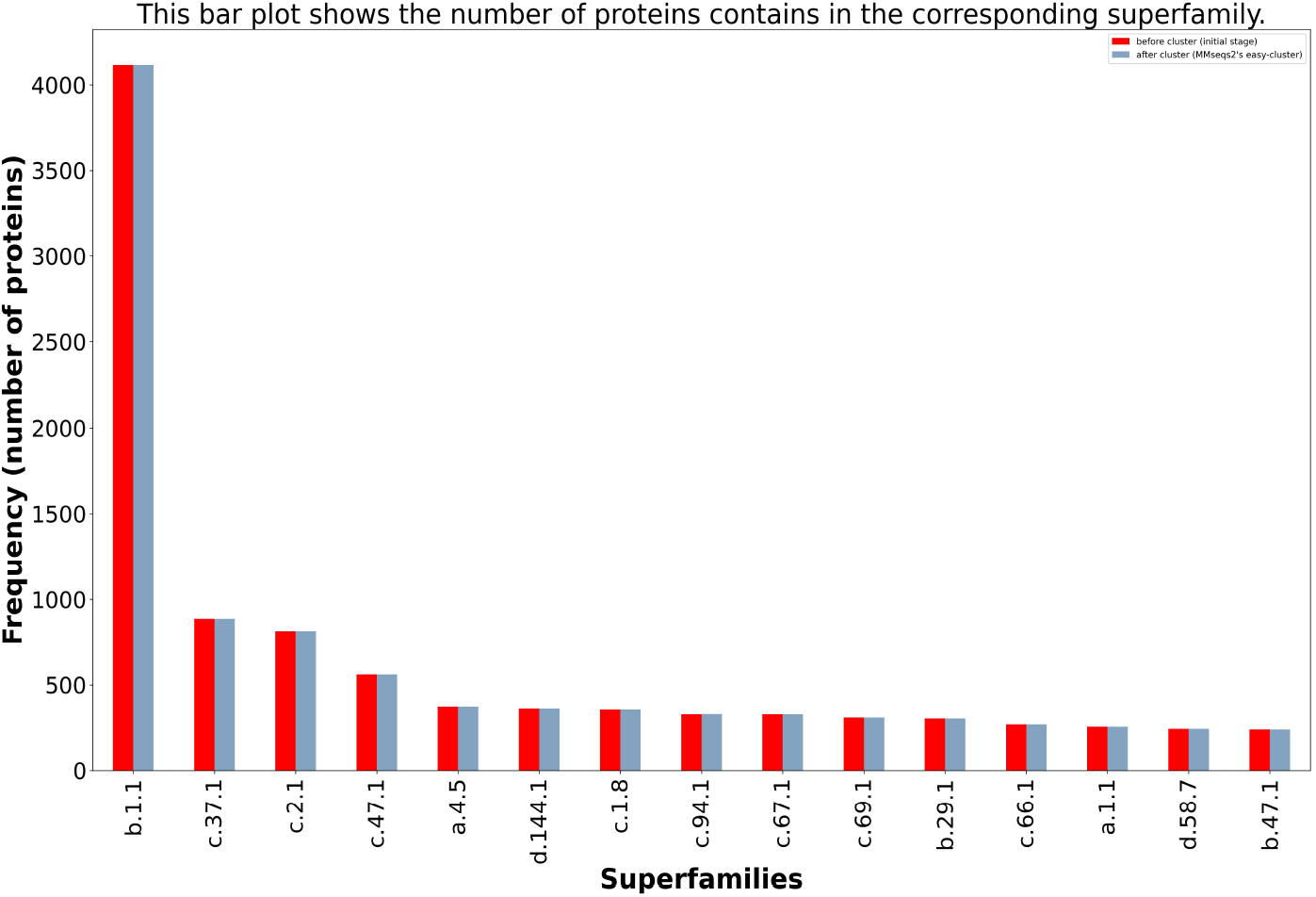
This image illustrates performance that is attained using MMseqs2 cluster algorithms. In this image, I particularly show the first 15 superfamilies, which contain maximum members.

The MMseqs2 is quite a remarkable tool for metagenomics and genomics sequence databases, as those fields deal with billions of sequences [4]. The strength and efficacy of the MMseqs2 clustering tools are proved in my experiment. For my dataset, which contains 24578 protein sequences, it takes fewer than ten seconds to cluster the dataset. The runtime of the `MMseqs2/Linclust` and `MMseqs2/easy-cluster` algorithms are almost linear, 𝒪 (*m ×N*) *≈* 𝒪 (*N*), which are not dependent on the number of clusters (K) ^17^. After scrutinizing the result, I assert that the conventional ML clustering techniques are inappropriate in protein clustering, as precision is highly recommended in genomics and proteomics research.

## 7 Discussion

After performing poorly in conventional ML clustering methods, I deliberately investigated the probable reason under the hood. The dissimilarities between traditional machine learning clustering and MMseqs2 clusters are enormous. The MMseqs2 tool produces clusters based on sequence identity and alignment coverage. In addition, those algorithms generate several cluster centroids based on similarity among the sequences. The number of centroids gets higher if the algorithm does not find many similarities and vice-versa. All in all, the MMseqs2’s clusters are deeply concerned regarding biological structures and their chemical orientations. Consequently, the tool uses several physicochemical properties’ information by utilizing BLOSUM, PAM, and VTLM metrics.

Although embeddings were performed based on protein functions, traditional ML algorithms exclusively relied on protein similarities (i.e., by employing cosine similarity metric). However, this information is inadequate; hence, biological information is required, like the MMseqs2 tool utilized.

## 8 Limitation of this experiment

First of all, because of limitations in computing power, I could not use the entire dataset in this evaluation. Secondly, the weakness of MMseqs2 is not investigated. Additionally, the MMseqs2 uses greedy approaches to build cluster centroids to speed up its process; it might sink into oblivion if it gets challenging datasets. I could have tested the tool using some challenging datasets to assure its reliability. Finally, I could have explored other widely used clustering and redundancy filtering tools, including UCLUST and CD-HIT ^18^, to justify the strength of MMseqs2 tool.

## 9 Challenges

I encountered various impediments while working on this project. First of all, the time and space complexities are NP-hard in KMeans and Hierarchical clustering algorithms. Due to this, the dataset was truncated to slightly mitigate the issue. Secondly, ESM pre-trained models contain millions of parameters; thus, it takes enormous time to embed the proteins. In my experiment, the ESM embedding models, ESM-1b and ESM-1v, took almost four days for protein embedding. Last but not least, a few days ago (August 22, 2022), an updated version of ESM was released (i.e., ESMFold or ESM2). The ESMFold is enormous; the largest model has 15 billion parameters. Consequently, it requires approximately 55 GB to store the uncompressed model weights, which is 4.5 times larger than the entire training set used (*UniRef50*).

## 10 Conclusion

In this paper, protein sequences are embedded using the ESM tool in order to execute conventional ML clustering methods, namely: KMeans and Hierarchical clusterings. Afterward, using the same data, two cluster techniques from the MMseqs2 tool are employed. These combinations of clusters are subsequently compared with each other.

In terms of protein sequence clustering, a rigorous analysis has been conducted among MMseqs2 clusterings and conventional ML clustering techniques and strategies. After a meticulous experiment, I recapitulate that MMseqs2 clustering methods are significantly better than conventional ML clustering algorithms. Several aspects of genomics and proteomics research, such as protein engineering and metagenomics, are directly associated with drug discovery. Therefore, we are required to use high-performing cluster methods to ensure the quality of clusters and data redundancy.

A hypothesis is made that the density-based clustering approaches could be conducive to protein clustering as enormous proteins have structural similarities and share a significant portion of the sequence with each other.

## Footnotes

1 From the header of protein sequences in the SCOPe database, it is easy to figure out the superfamily (alphabeti-cal.numerical.numerical) and family (alphabetical.numerical.numerical.numerical) patterns.

2 ASTRAL Protein Datasets: https://scop.berkeley.edu/astral/subsets/ver=2.08

3 https://github.com/mrzResearchArena/protein-clustering/blob/main/choose-K-clusters.ipynb

4 https://github.com/mrzResearchArena/protein-clustering/blob/main/species-analysis.ipynb

5 MAFFT tool: https://academic.oup.com/nar/article/30/14/3059/2904316/

6 Here, N, K, I, and D represent the number of protein sequences, number of clusters, number of iterations, and dimension of the dataset, respectively.

7 https://github.com/mrzResearchArena/protein-clustering/blob/main/choose-K-clusters.ipynb

8 https://github.com/mrzResearchArena/protein-clustering/blob/main/choose-K-clusters.ipynb

9 https://github.com/mrzResearchArena/protein-clustering/blob/main/ESM-embedding.ipynb

10 https://github.com/mrzResearchArena/protein-clustering/blob/main/ESM-embedding.ipynb

11 Although ESM-1v has five different variant, I chose only the first one (i.e., **esm1v_t33_650M_UR90S_1**, esm1v_t33_650M_UR90S_2, esm1v_t33_650M_UR90S_3, esm1v_t33_650M_UR90S_4, esm1v_t33_650M_UR90S_5).

12 https://github.com/mrzResearchArena/protein-clustering/blob/main/Clustering-kmeans-HC.ipynb

13 https://github.com/mrzResearchArena/protein-clustering/blob/main/lookupTable.ipynb

14 https://github.com/mrzResearchArena/protein-clustering/blob/main/MMseqs2-Clustering.ipynb

15 https://github.com/mrzResearchArena/protein-clustering/blob/main/MMseqs2-Clustering.ipynb

16 https://github.com/mrzResearchArena/protein-clustering/blob/main/MMseqs2-Clustering.ipynb

17 **m** denotes the number of k-mers. It is usually 20-25 for protein sequences. Additionally, **N** indicates the number of sequences.

18 http://weizhong-lab.ucsd.edu/cdhit_suite/

